# Insights into the biosynthesis of icumazole unveiling a distinctive family of crotonyl-CoA carboxylase/reductase

**DOI:** 10.1101/2022.11.22.517467

**Authors:** Feng Xie, Alexander F. Kiefer, Anna K. H. Hirsch, Olga Kalinina, Chengzhang Fu, Rolf Müller

**Affiliations:** Helmholtz Institute for Pharmaceutical Research Saarland (HIPS), Helmholtz Centre for Infection Research (HZI), and Department of Pharmacy, Saarland University, 66123 Saarbrücken, Germany; Helmholtz International Lab for Anti-Infectives, Helmholtz Center for Infection Research, 38124 Braunschweig, Germany; German Centre for Infection Research (DZIF), 38124 Braunschweig, Germany; Medical Faculty, Saarland University, 66421 Homburg, Germany; Center for Bioinformatics, Saarland Informatics Campus, 66123 Saarbrücken, Germany

**Author notes:** Corresponding authors: Rolf Müller, Chengzhang Fu. Lead Contact: Rolf Müller.

**Keywords:** Crotonyl-CoA carboxylase/reductase, icumazole, thuggacin, myxobacteria, polyketide synthase, biosynthesis, sequence similarity network

## Abstract

Icumazoles are potent antifungal polyketides with intriguing structural features. Here, we present the polyketide synthase (PKS)/nonribosomal peptide synthetase (NRPS) hybrid biosynthetic gene cluster of icumazoles. Surprisingly, an unusual non-terminal thioesterase domain divides the PKS/NRPS assembly line. The succeeding PKS modules potentially form a rare precursor 4-methyl-2-hexenoyl-ACP thus deviating from the previously proposed polyoxypeptin pathway. The 4-methyl-2-hexenoyl-ACP is further reductive carboxylated to 2-methylbutylmalonyl-ACP essential for icumazole biosynthesis by IcuL, representing a new type of crotonyl-CoA carboxylase/reductase (CCR). We characterize IcuL and its homologs TgaD and Leu10 *in vitro*, suggesting a stricter substrate specificity of this new family of CCRs than found in canonical ones. Intriguingly, we also find that TgaD unprecedently utilizes both NADPH and NADH as cofactors with similar efficiency, diverging from the NADPH-specific characteristic of canonical CCRs. Furthermore, a sequence similarity network-based bioinformatic survey reveals that the IcuL-like CCRs are evolutionarily separated from canonical CCRs.

## Introduction

Natural products are one of the most important sources for human and veterinary drug discovery as well as agrochemicals (Newman and Cragg, 2020). Gram-negative myxobacteria are prolific producers of novel bioactive small molecules. Numerous secondary metabolites with various bioactivities have been characterized (Weissman and Müller, 2010; Herrmann et al., 2017), such as anticancer drug epothilones (Bollag et al., 1995; Gerth et al., 1996), broad-spectrum antibacterials cystobactamids (Baumann et al., 2014; Hüttel et al., 2017), HIV inhibitors aetheramides (Plaza et al., 2012), C1 metabolism-targeting biofilm inhibitor carolacton (Jansen et al., 2010; Fu et al., 2017), and antifungal icumazoles (Barbier et al., 2012).

Icumazole A (**1**) and its derivatives B1 (**2**) and B2 (**3**) featuring isochromanone, triene, and oxazole moieties were isolated from *Sorangium cellulosum* So ce701 (Barbier et al., 2012). However, the biosynthesis of icumazoles remained elusive. Icumazoles are polyketides containing an oxazole ring, as often observed in microbial natural products produced by hybrid biosynthetic gene clusters (BGCs) of polyketide synthases (PKSs) and nonribosomal peptides synthetases (NRPSs). Type I PKS and NRPS are multi-domain megaenzymes organized into catalytic modules (Fischbach and Walsh, 2006). Each module consists of multiple catalytic domains responsible for incorporating one building block to extend and modify the nascent intermediates tethered to the biosynthetic enzymes via acyl/peptidyl carrier protein (ACP/PCP) domains. Closer scrutiny of the chemical structures of icumazoles suggested that the biosynthesis of their 2-methylbutane side chain requires an unusual 2-(*S*)-2-(2-methylbutyl)malonyl-CoA (MBM-CoA) as a PKS building block (Figure 1A). To date, two different biosynthetic pathways of α-alkylmalonyl-CoAs are known: the reductive carboxylation of α,β-unsaturated acyl-CoAs by crotonyl carboxylase/reductases (CCRs) (Wilson and Moore, 2012) and the direct carboxylation of acyl-CoAs by acyl-CoA carboxylases (YCCs) (Ray et al., 2016). Additionally, alcohol dehydrogenases (ADHs) with very low sequence similarity to the well-studied CCRs were proposed for the formation of the corresponding alkylmalonyl-CoAs in pathways of myxobacterial compounds leupyrrin A_1_ (**4**) (Kopp et al., 2011) and thuggacin A (**5**) (Buntin et al., 2010a) (Figure 1A).

**Figure 1.**
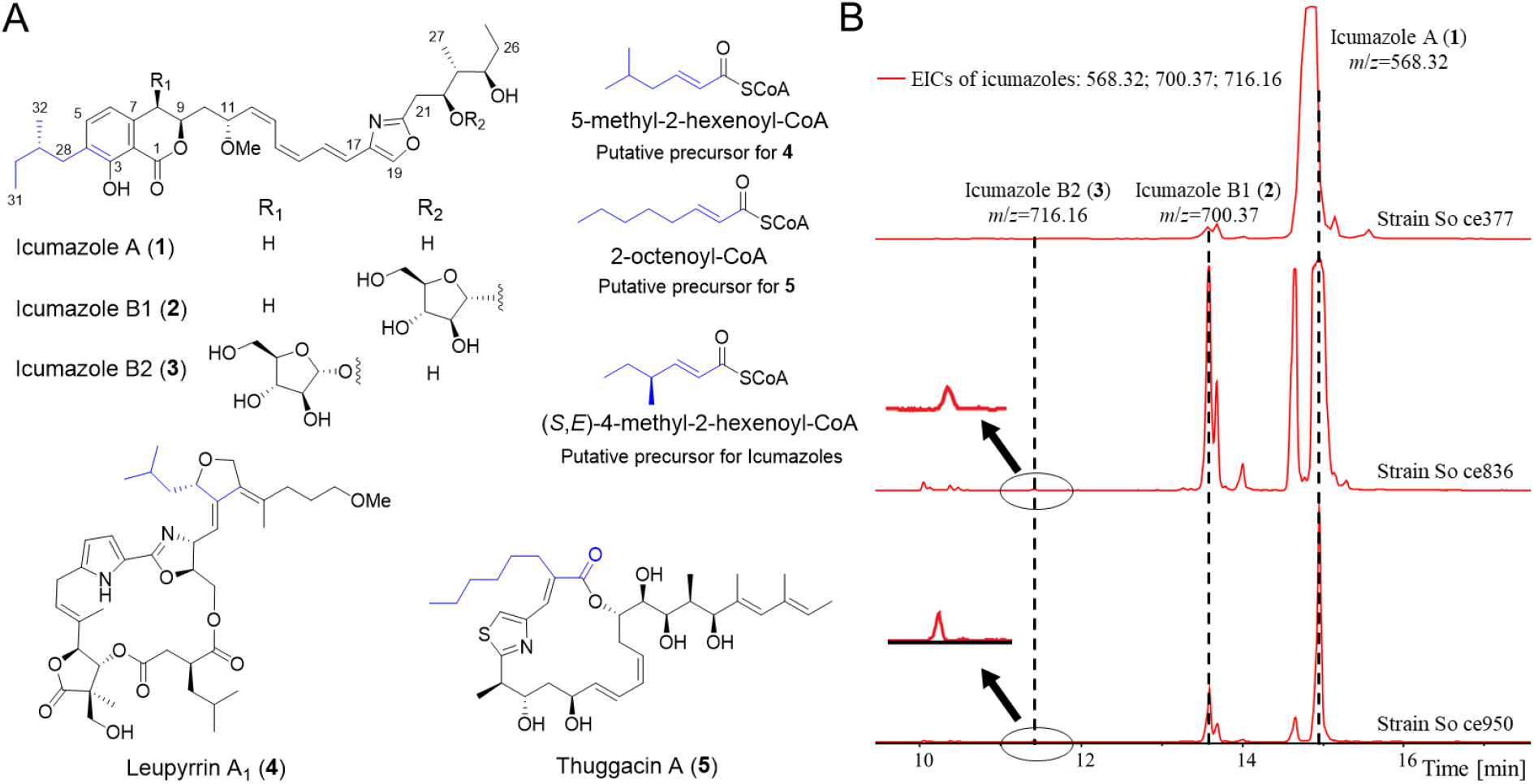
Myxobacterial compounds with side chains introduced by noncanonical CCRs. (A) Representative compounds structures and their putative precursors. The carbon atoms derived from α-alkylmalonyl-CoAs are marked in blue. (B) Production of icumazoles in three *Sorangium* strains. Extracted ion chromatograms (EICs) of icumazoles from respective strains.

In this work, we discovered the icumazole BGC and proposed a biosynthetic pathway of icumazoles, which includes a PKS/NRPS hybrid assembly line and a CCR-based route to produce the precursor MBM-ACP. Further, *in silico* analysis of a CCR from this BGC, IcuL, unveiled a noncanonical class of CCRs, which was found to exhibit significantly stricter substrate specificity distinguishing it from biochemically earlier characterized CCRs. One member from the IcuL-like CCR family, TgaD, stands out due to its unique ability to utilize NADH in contrast to the strict NADPH specificity of previously biochemically characterized CCRs (Quade et al., 2011).

## Result and Discussion

### Identification of the icumazole BGC

Three *S. cellulosum* strains from our in-house myxobacterial strain collection exhibited icumazole production capability, including two strains with available genome data (complete genome sequence So ce836 and a draft genome for So ce377; Figure 1B). The BGC prediction in the So ce836 genome (Table S1) led to the identification of only one large modular PKS/NRPS BGC (BGC19) that has one analog in the genome of So ce377 (Table S2).

More hints towards the correct identification of the candidate BGC came from the correlation of the gene functions with the structural features of icumazoles. First, the isochromanone ring should to be assembled by a PKS and a unique TE domain based on the reported biosynthesis of ajudazol (Buntin et al., 2010b). Second, we assumed the 2-methylbutyl side chain of the isochromanone to derive from MBM-CoA that was formed by a CCR as found encoded in the BGC. Third, the presence of an oxazole moiety in icumazoles indicates the requirement of a NRPS module featuring an adenylation (A) domain that activates serine, a heterocyclization (HC) domain for the oxazoline ring formation, and an oxidation (OX) domain to eventually form the oxazole (Buntin et al., 2010b). Furthermore, the oxazole module is supposed to be the only NRPS module, and three PKS modules should precede the oxazole module according to the so-called co-linearity rule (Fischbach and Walsh, 2006). All these features are consistent with the gene function analysis of BGC19, consisting of 14 PKS modules and 1 NRPS module encoded by 10 PKS/NRPS genes and 1 noncanonical CCR (Table S2). Moreover, a transcriptome study of So ce836 showed that *icuA-L* were transcribed coincidently with the production of icumazoles and also suggested the boundary of the *icu* BGC (unpublished results). Consequently, BGC19 was considered the icumazole BGC (designated as *icu*).

### Biosynthetic pathway of icumazoles

The *icu* BGC comprises more PKS modules than needed for building the carbon skeleton, indicating that the additional PKS modules are inactive or responsible for some noncanonical part of the carbon backbone (Figure 2A, Table S2). To determine the potential function of each domain, they were analyzed for the known critical residues (Data S1), including those for acyltransferase (AT), ketosynthase (KS), ketoreductase (KR), dehydratase (DH), enoylreductase (ER), ACP and thioesterase (TE). The shorter AT_8_ and AT_14_ lack conserved catalytic residues, indicating their possible disability of loading building blocks to their cognate ACPs. KS_12_ has a C to F point mutation in the catalytic site, similar to the C to Q mutation in KS_Q_ (Chisuga et al., 2022), suggesting loss of condensation function. We also observed potential nonfunctional optional domains, including DH_10_, DH_11_, DH_14_, and ER_13_, which do not affect the PKS chain extension by the corresponding modules.

**Figure 2.**
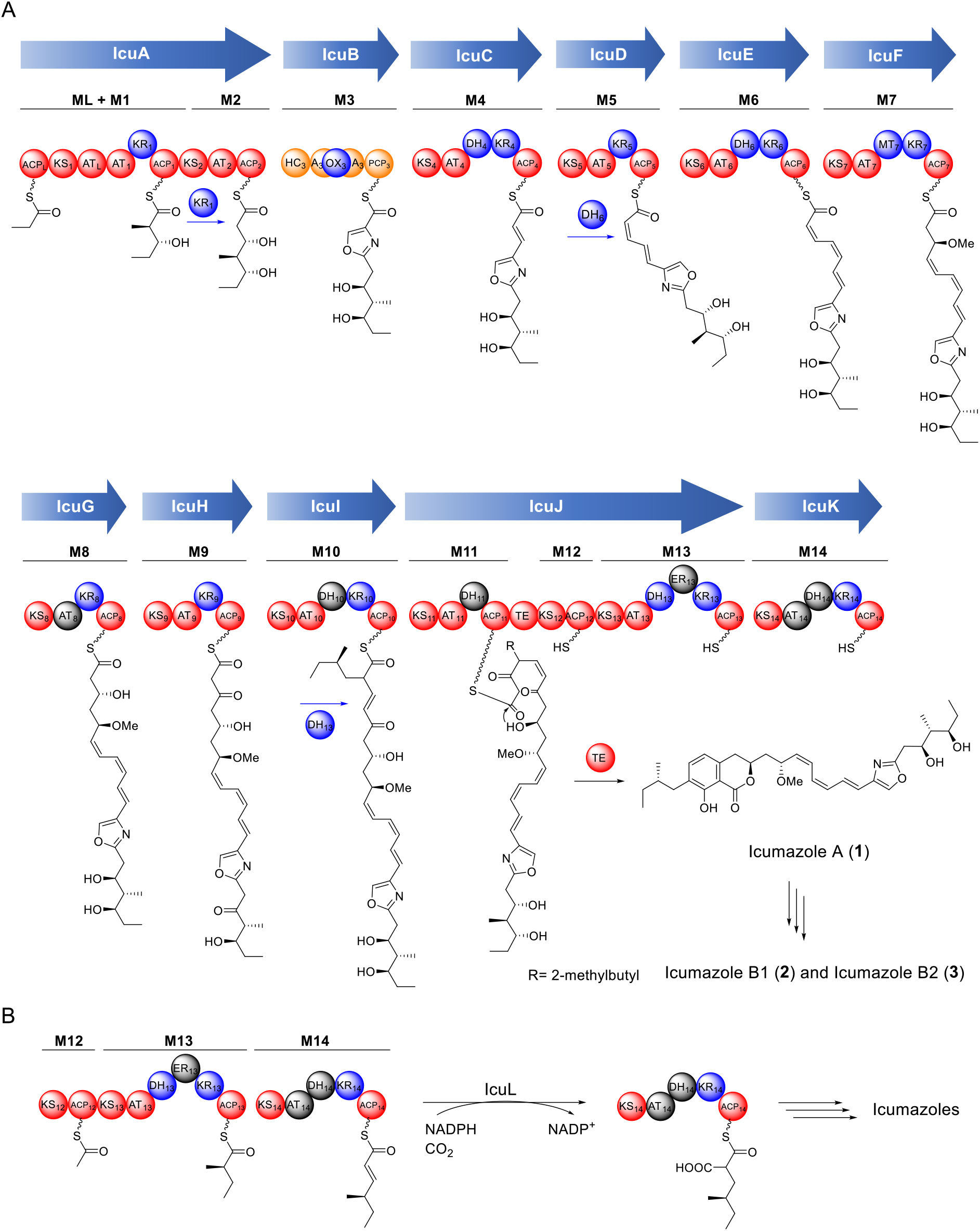
Proposed biosynthetic pathway of icumazoles. (A) Scheme of the PKS/NRPS assembly line of icumazoles. Functional domains of PKS and NRPS are shown in red and orange, respectively; The optional domains are shown in blue; Proposed inactive domains are shown in gray. Blue arrows indicate possible cross-modular catalyses. For details on the abbreviations, see the main text. For the sequence alignment analysis of each domain, see Data S1. (B) Proposed pathway of the precursor 2-methylbutyrylmalonyl-ACP in the icumazole biosynthesis.

The icumazole megasynthetase adopts a unique architecture as found in other myxobacterial PKS, where an additional AT domain (AT_L_) is embedded in the first extending module, forming a mixed module exhibiting ACP_L_-KS_1_-AT_L_-AT_1_-KR_1_-ACP_1_ organization (ML+M1, Figure 2A). We proposed that AT_L_ activated and transferred the starter unit onto ACP_L_, in line with the reported biosynthesis of **5** (Buntin et al., 2010a), ajudazol (Buntin et al., 2010b), and stigmatellin (Gaitatzis et al., 2002). However, unlike in other cases, AT_L_ from the *icu* pathway was predicted to recognize methylmalonyl-CoA as substrate (Table S2), which is supported by the critical R residue for selecting the dicarboxylic units in AT_L_ (Data S1A) (Rangan and Smith, 1997; Long et al., 2002; Liou et al., 2003). KS_1_ was proposed to bifunctionally catalyze the KSQ-like decarboxylation in the loading step and the condensation in the first extension step. The KR_1_ was a B-type KR that generates a D-configuration hydroxyl group according to the sequence alignment (Data S1C), which is in agreement with the *R*-configuration at C24 in icumazoles as confirmed by total synthesis (Figure 1A) (Barbier et al., 2012). The second extending module (M2) lacks a KR domain, which is inconsistent with the corresponding hydroxyl group at C22 of icumazoles. We hypothesized that the ketoreduction in M2 may be catalyzed *in-trans* by KR_1_, while a similar case was observed in the azalomycin biosynthesis where a *cis* ER *in-trans* is assumed to catalyze the enoyl reduction in a neighboring module (Zhai et al., 2020). Module 3 consists of a HC domain, an A domain, an OX domain and a PCP domain (Figure 2A), forming a typical oxazole or thiazole biosynthetic module as observed in ajudazol (Buntin et al., 2010b) and epothilone (Dowling et al., 2016) biosynthesis. The following modules, M4 to M6, are assumed to further extend the PKS chain giving rise to the triene moiety in icumazoles. However, these three modules only have two active DHs (Table S2 and Data S1D), DH_4_ and DH_6_, implying that one of them might act twice during the biosynthesis. The dehydration in M5 that lacks a DH domain might be cross-modularly catalyzed by DH_6_ because the double bonds supposed to be formed by M5 and M6 are both in *Z*-configuration. The presence of KR_7_ and MT_7_ in module 7 is in line with the *O*-methyl group at C11 in icumazoles.

Subsequently, the *icu* biosynthesis enters the formation of the isochromanone ring resembling the TE-mediated cyclization mechanism in ajudazol biosynthesis (Buntin et al., 2010b). Unlike the terminal TE from the ajudazol pathway, the icumazole TE is situated between PKS modules 11 and 12 in IcuJ. A PKS or NRPS with a non-terminal TE is highly unusual (Little and Hertweck, 2022). Moreover, the actual function of this kind of TEs is still unclear except for the TE of Fr9C catalyzing the formation of a *cis*-double bond in FR901464 biosynthesis (He et al., 2014) and the TE-B enabling chain release in the malleicyprol pathway (Little et al., 2022). The TE domain in IcuJ possesses the conserved catalytic triad (Data S1G), suggesting its role in releasing **1**. This function was further supported by phylogenetic analysis in which the icumazole TE clustered with the ajudazol TE (Data S1H). Intriguingly, modules M8 to M11 located before the TE seem almost collinear with PKS chain extension required before isochromanone formation. Although AT_8_ seems inactive, M8 may still be functional, aided by an adjacent AT acting *in-trans*. An alternative would be that M8 is dysfunctional, whereas M9 works iteratively in this pathway. M10 is likely to elongate the nascent PKS backbone, utilizing the unique MBM building block (see below). Module 11 finalizes chain extension by using a methylmalonyl-CoA extender prior to chain release/ring formation by the TE (Figure 2A). The inactive DH_10_ and DH_11_ (Data S1D) indicate the involvement of a DH acting *in-trans* in M10. It is thus proposed that *cis*-domains act *in-trans* multiple times in icumazole biosynthesis. However, the exact mechanism is yet to be characterized. After the chain release and cyclization, remote unspecific glycosyltransferases are expected to mediate the glycosylation of **1** because no glycosyltransferase gene is found nearby the *icu* BGC (Figure 1B).

### An unusual modular PKS-dependent biosynthesis of the MBM moiety

The activated MBM is a scarce PKS building block that was only reported once in the biosynthetic pathway of polyoxypeptin A (Umezawa et al., 2001). A feeding study suggested that the MBM moiety in polyoxypeptin A is derived from isoleucine. A follow-up *in silico* analysis of the polyoxypeptin A BGC proposed a biosynthetic pathway to MBM, starting from isoleucine to form the corresponding α-keto acid (Du et al., 2014). The α-keto acid was thought to be extended by a FabH-like KS while tethered to a discrete ACP. Genes encoding α-keto acid dehydrogenases were also proposed to be involved in the following steps for producing the (*S,E*)-4-methyl-2-hexenoyl-CoA (MHE-CoA) as the substrate for the corresponding CCR. However, no genes encoding such enzymes could be discovered nearby the *icu* BGC except for the proposed CCR-encoding gene *icuL*. Moreover, these genes do not form an operon located elsewhere in the So ce836 genome. Thus, all available evidence suggests a distinctive biosynthetic route to form MBM for icumazole biosynthesis.

As discussed above, three PKS modules of unassigned function follow the unusual non-terminal TE domain in IcuJK. Modules 12 and 13 were found in IcuJ, with M11 at the *N*-terminus and the TE domain located before M12. IcuK comprising M14 is a single-module protein. We hypothesized that ACP_12_ was self-malonylated and decarboxylated by the KS_Q_-like KS_12_. M13 next extends the acetyl-*S*-ACP species with one methylmalonate and reduces the β-keto to form a 2-methylbutyryl moiety attached to ACP_13_. The nascent chain is further elongated by M14 with one malonyl-CoA (Figure 2B) and subsequently KR_14_ and DH_14_ reduces the β-keto species to form a MHE moiety tethered to ACP_14_, exactly representing the required substrate for a CCR reaction forming the extender unit required for icumazole biosynthesis, albeit in the ACP-bound form instead of the common CoA substrate for CCRs. According to the sequence alignment, ER_13_, AT_14_, and DH_14_ were predicted inactive due to the loss or mutation of key residues (Data S1). However, myxobacterial natural products’ PKS and NRPS biosynthesis often deviate from the textbook rules (Keatinge-Clay, 2017) and *in-trans* acting domains could supplement the missing function as proposed for other parts of the icumazole biosynthesis.

### IcuL represents a unique group of CCR

The only non-PKS/NRPS gene in *icu* BGC is *icuL* and its encoded protein shows high identity to TgaD from the **5** pathway (48.6%) and Leu10 from the **4** pathway (52.6%). TgaD and Leu10 were proposed to be CCRs, albeit exhibiting extremely low similarity to the canonical CCRs (Buntin et al., 2010a; Kopp et al., 2011). Similarly, IcuL only exhibits 18% identity to CinF (Quade et al., 2011), a well-characterized actinobacterial CCR. Also, the ~380 aa average length of IcuL-type CCR is roughly 70 aa shorter than CinF-type CCR which is ~450 aa. Most importantly, the actual function of IcuL, TgaD, and Leu10 are lacking experimental evidence (Buntin et al., 2010a; Kopp et al., 2011; Wilson and Moore, 2012). We here propose that IcuL is involved in icumazole biosynthesis by providing MBM-ACP from MHE-ACP by reductive carboxylation (Figure 2B). To verify this hypothesis and to investigate the characteristics of this new type of CCRs, we overexpressed and purified IcuL, TgaD, and Leu10 as recombinant proteins. Six *N*-acetylcysteamine (SNAC) thioester mimicking substrates with different chain lengths, namely 4-methyl-2-pentenoyl-SNAC (**S1**), 5-methyl-2-hexenoyl-SNAC (**S2**), (*S*,*E*)-4-methyl-2-hexenoyl-SNAC (**S3**), 2-pentenoyl-SNAC (**S4**), 2-hexenoyl-SNAC (**S5**), and 2-octenoyl-SNAC (**S6**), were chemically synthesized (Figure 3A, Data S2) to characterize these enzymes *in vitro*.

**Figure 3.**
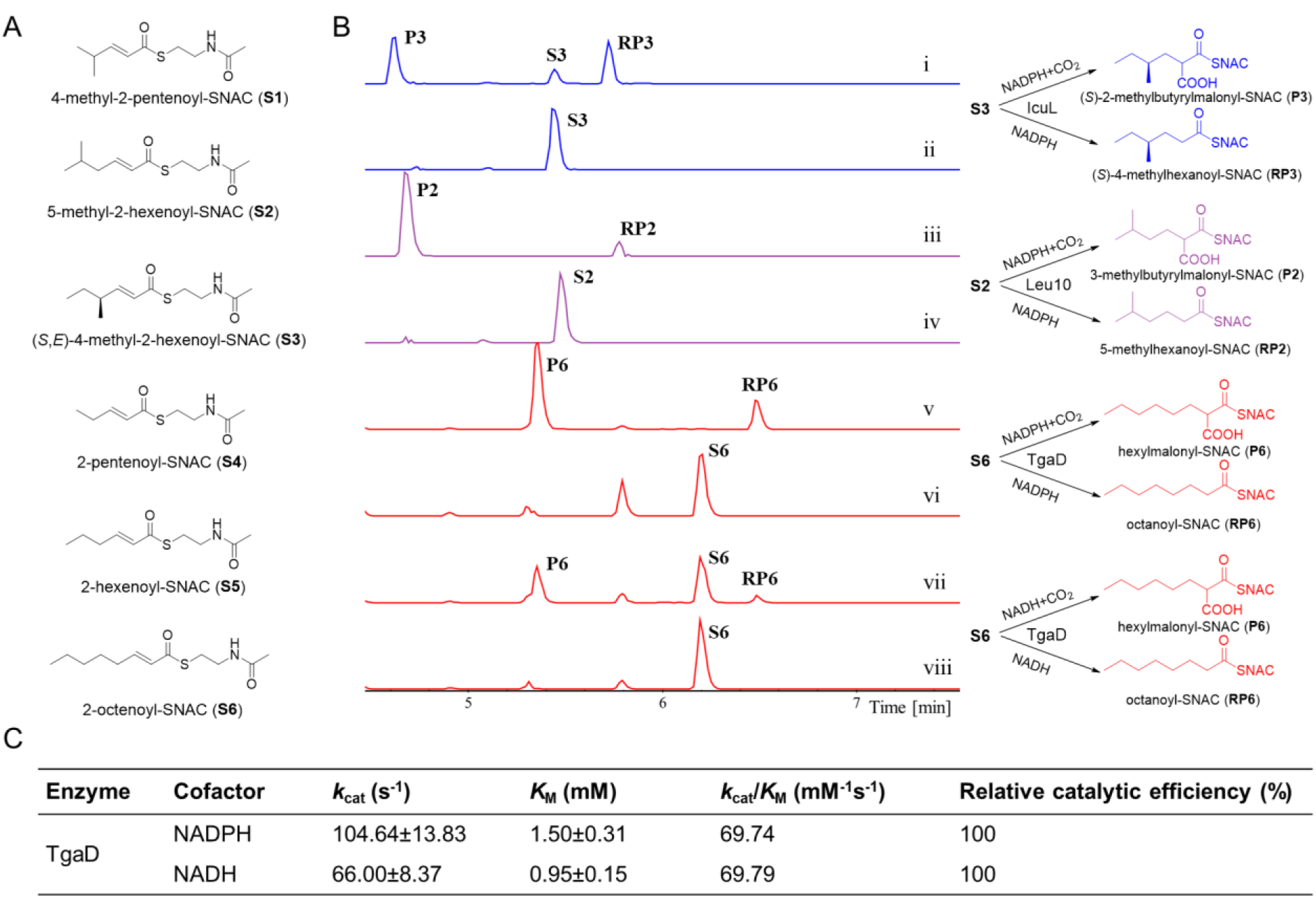
Biochemical studies of IcuL-like CCRs. (A) Structure of the synthetic substrates. (B) *In vitro* assays of IcuL (i and ii), Leu10 (iii and iv) and TgaD (v to viii). EICs of substrates and the corresponding products are shown in the left panel. Illustration of enzymatic reaction of each compound is shown in the right panel. Traces i, iii, v and vii show reactions containing enzymes, cofactors, NaHCO_3_ and corresponding substrates; traces ii, iv, vi and viii are control reactions without enzymes. Traces i to vi use cofactor NADPH, while traces vii and viii use NADH. EICs of i to iv = 230.12; 232.14; 276.13; EICs of v to viii = 244.14; 246.15; 290.14. **P2**: reductive carboxylated product of **S2** (276.13); **RP2**: reduced product of **S2** (232.14); **P3**: reductive carboxylated product of **S3** (276.13); **RP3**: reduced product of **S3** (232.14); **P6**: reductive carboxylated product of **S6** (290.14); **RP6**: reduced product of **S6** (246.15). See also Figure S1. (C) Enzyme kinetic analysis of TgaD towards NADH/NADPH. See also Figure S2.

We employed *icuL* genes from different strains, but only IcuL from So ce377 was soluble expressed and purified with a *N*-terminal MBP tag. After an overnight incubation of IcuL with reduced nicotinamide adenine dinucleotide phosphate (NADPH), NaHCO_3_ and SNAC-ester **S3**, two new products with *m/z* of 232.14 and 276.13 were formed (Figure 3B, i and ii), which are 2 dalton and 46 dalton heavier than the *m/z* value of **S3** (230.12), respectively, suggesting the formation of a reduced product 4-methylhexanoyl-SNAC (**RP3**) and a reductive carboxylated product MBM-SNAC (**P3**). The same types of products were also detected in assays of Leu10 (Figure 3B, iii and iv) and TgaD (Figure 3B, v and vi), corroborating their function as CCRs.

We then tested the substrate specificity of these three CCRs using all six alkenoyl-SNACs. IcuL catalyzed the reductive carboxylation of **S2** and **S5** at a moderate level compared to **S3** with the authentic chain length, while almost no conversion was observed when using **S1**, **S4** and **S6** as substrates (Figure S1A). Except for the authentic SNAC **S2**, Leu10 is active employing **S5** while it was found almost inactive on other SNAC-esters (Figure S1B). We examined the structures of these substrates and found that **S2**, **S3** and **S5** share the same carbon chain length (Figure 3A), suggesting a crucial role of chain length in the substrate specificity of this new type of CCRs. In line with this assumption, none of the remaining five substrates, at least two carbons shorter than **S6**, can serve as substrate for TgaD (Figure S1C). This property is divergent from CinF from the cinnabaramide BGC (Quade et al., 2011) and PteB from the filipin BGC (Yoo et al., 2011), which are comparably much more promiscuous towards substrates with various carbon chain lengths. The CCR substrate specificity is further supported by the fact that the IcuL-like CCRs only introduce one kind of side chain in icumazoles, thuggacins, and leupyrrins, respectively.

### TgaD utilizes both NADH and NADPH as cofactor

CCRs are recognized to be NADPH-dependent enzymes (Wilson and Moore, 2012). Quade et al. determined that CinF strictly uses NADPH as its cofactor based on the *in vitro* biochemical characterization and co-crystal structures (Quade et al., 2011), presumably implying the general feature of tight cofactor specificity in canonical CCRs. We asked the question if members of the newly identified CCR family are capable of leveraging NADH as a cofactor. As expected, the catalytic activities of both, IcuL and Leu10, were abolished when employing NADH as the cofactor (Figure S1D). However, octanoyl-SNAC (**RP6**) and hexylmalonyl-SNAC (**P6**) were successfully produced from **S6** by TgaD when using NADH instead of NADPH (Figure 3B, vii and viii).

To assess the specificity of TgaD towards NADH and NADPH, we decided to measure the kinetic parameters of TgaD. However, all six alkenoyl-SNACs degraded quickly (Figures S1A-1C, S2A-2B), affecting an accurate kinetic study. In order to identify the reason for the degradation, we incubated **S6** with each component from the reaction separately. The degradation only appeared when Tris-HCl 8.0 and NaHCO_3_ (Figure S2C) were used, suggesting that the relatively high pH should be the leading cause of the degradation. Therefore, a suitable condition for the CCR kinetic assay employing the SNAC substrates was required. Considering the pH of NaHCO_3_ (8.3), we first tested the TgaD assay in various buffers without bicarbonate (Figure S2D) revealing that the degradation of **S6** slowed down when using buffers with pH lower than 7.0. Further experiments showed that adding NaHCO_3_ promotes degradation intensely even at a concentration of 25 mM when using pH 7.0. The degradation took place similarly when using NH_4_HCO_3_ at pH 7.8. Buffers with pH lower than 7.0 are considered inappropriate for TgaD catalysis due to the low velocity. It is noteworthy that the turnover number of TgaD was only determined by the enzyme and NAD(P)H, regardless of the presence/absence of bicarbonate. Taken together, the Tris-HCl 7.0-based reaction system without bicarbonate seems the ideal condition for kinetic analysis of cofactors NADH and NADPH. By comparison of *k*_cat_ and *K*_M_ constants, we found that TgaD processes these two cofactors with similar efficiency (Figures 3C and S2E). The *k*_cat_ value of TgaD using NADPH is 1.5-fold higher than employing NADH, indicating a higher catalytic velocity of NADPH than NADH. Unexpectedly, TgaD has a higher *K*_M_ value when using NADPH, suggesting a higher binding affinity for NADH than for NADPH. However, TgaD generally showed the same level of preference for both NADH and NADPH. To the best of our knowledge, TgaD is the first CCR reported to utilize both NADH and NADPH.

CCRs play a vital role in highly energy-consuming natural product biosynthesis by supplying special building blocks (Wilson and Moore, 2012). The NADH-utilizing property of CCR provides an opportunity to engineer the metabolic flux for production improvement since the generation of NADP(H) is performed by NAD kinase from NAD(H) in bacteria (Spaans et al., 2015), involving consumption of ATP. A recent study reported establishing a CO_2_ fixation synthetic cycle, where two CCRs play a core role in the pathway (Schwander et al., 2016). Our finding thus enables future efforts on alteration of the cofactor preference of these CCRs, which may result in more efficient *in vivo* pathways by saving two molecules of ATP for each catalytic cycle.

### Distribution of IcuL/TgaD-like CCRs in nature

To systematically study this new CCR family, we conducted a global sequence-based survey from the UniProt database (The UniProt Consortium., 2021) as described in STAR Methods (Figure 4). As expected, IcuL, Leu10, and TgaD cluster in a unique group with eight other proteins (Table S3) apart from the canonical CCRs (a cluster of 586 proteins, including CinF (Quade et al., 2011), AntE (Yan et al., 2012), SalG (Eustáquio et al., 2009), TcsC (Blažič et al., 2015) and PteB (Yoo et al., 2011), see insets in Figure 4), suggesting their potential functional difference. Surprisingly, we also found that AsmB6 (UniProt ID A0A5P2GIQ0) which was proposed as the CCR for ansaseomycin biosynthesis was grouped into the new IcuL-like CCR family (Liu et al., 2019). Moreover, the CCR from polyoxypeptin A (Du et al., 2014) (identity/similarity of 41.6%/58.6% to IcuL) should also belong to this IcuL-like family but was absent because the corresponding genome data was missing in the public databases. Closer scrutiny of the structures of icumazoles, leupyrrins, thuggacins, ansaseomycins and polyoxypeptins, which are all known compounds using the new family CCR in biosynthesis, revealed that only one kind of side chain was found (Umezawa et al., 1998; Bode et al., 2003; Irschik et al., 2007; Steinmetz et al., 2007; Barbier et al., 2012; Liu et al., 2019), suggesting a precise substrate specificity of the IcuL-like CCRs controlling the chain length of building blocks for the polyketide chain extension. Detailed mechanistic studies are thus required to fully elucidate the biochemistry of this new subclass of enzymes.

**Figure 4.**
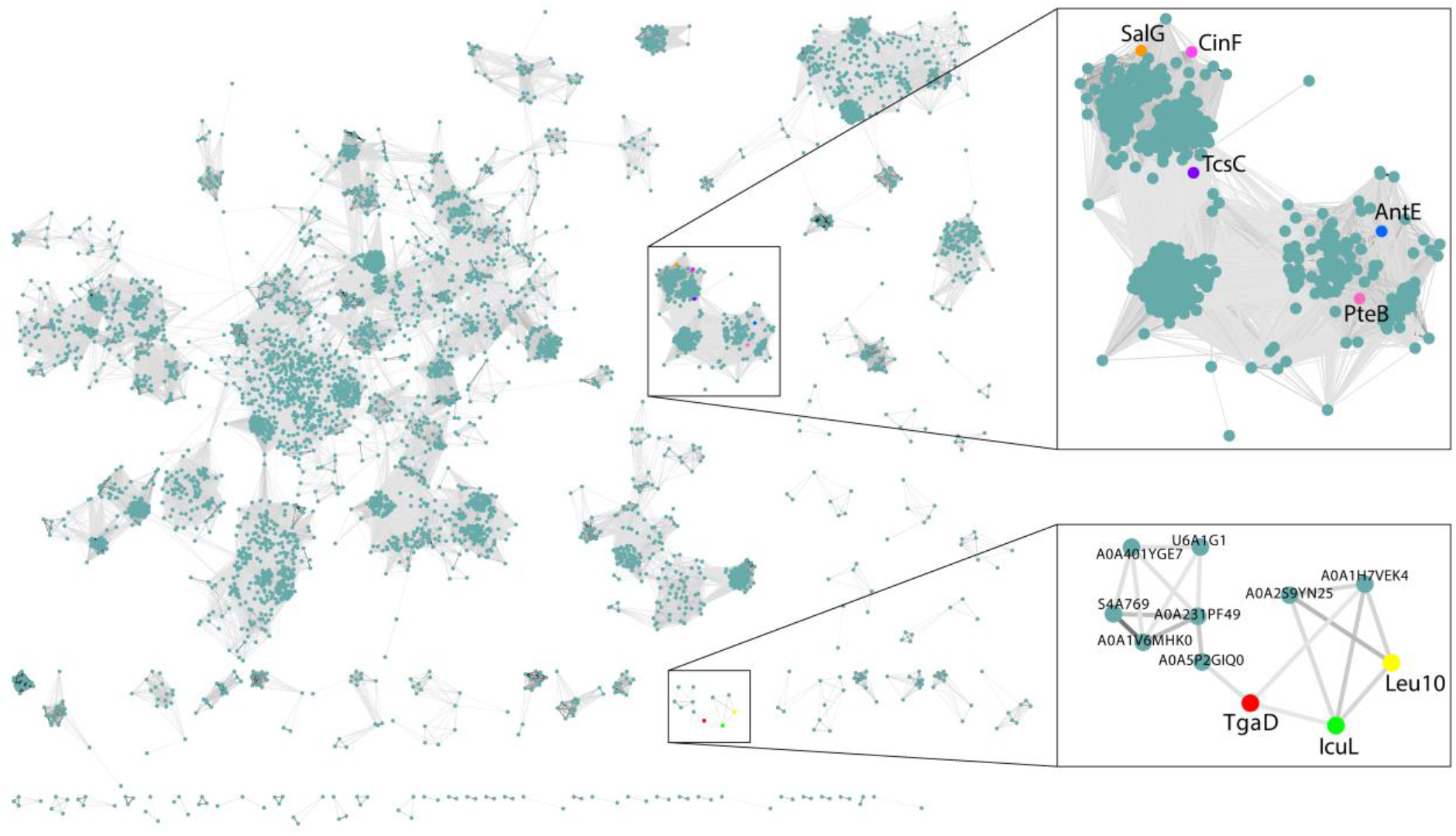
Global distribution of ADH family proteins located within BGCs. The canonical CCR family (upper inset) and the new CCR family are highlighted (underneath). The edges are drawn between nodes, representing individual ADHs (darker edges indicate >25% sequence identity values). CCRs from well characterized BGCs are colored and labeled.

### Significance

The myxobacterial polyketides icumazoles exhibit potent antifungal bioactivity. However, the biosynthetic pathway of icumazoles remained unknown, leaving an especially intriguing open question of how the 2-methylbutyryl side chain of the isochromanone ring is introduced. Here, we describe the BGC of icumazoles, a hybrid pathway consisting of ten PKS genes, one NRPS gene, and one CCR gene (*icuL*). It is worth noting that the PKS/NRPS assembly line is divided into two parts by an unusual non-terminal TE domain. The former part comprises a loading module and 11 extension modules, which are proposed to build the backbone of icumazoles while requiring a highly unusual MBM building block. The latter PKS modules might be responsible for forming a rare precursor MHE-ACP, which deviates from the previously proposed MHE pathway. MHE-ACP is further converted to MBM-ACP by IcuL as a PKS building block for icumazole biosynthesis. We studied IcuL and its homologs TgaD and Leu10 *in vitro* revealing a stricter substrate specificity of the IcuL-like CCRs compared to canonical ones. Most intriguingly, we showed that TgaD unprecedently utilizes both NADPH and NADH as the cofactor with similar efficiency, diverging from the NADPH-specific characteristic of canonical CCRs. Furthermore, a global sequence survey shows that the IcuL-like CCRs cluster is a new group separated from the canonical CinF-like CCRs unambiguously. Further in-depth studies of the IcuL-like CCR, especially TgaD, are promising for synthetic biology applications because of their strict substrate specificity and the distinctive NADH-utilizing characteristic of TgaD, which will contribute to synthetic carbon fixation and production of high-value-added chemical products.

## Acknowledgments

This work was supported by grants to R.M. and C.F. from the following sources: Deutsche Forschungsgemeinschaft and the Helmholtz International Lab.

## Author Contributions

F.X., C.F., and R.M performed the gene cluster analysis. F.X. and C.F. performed and analyzed biochemical experiments. A.F.K. and A.K.H.H. synthesized the SNAC substrates. O.K. and C.F. performed and interpreted the SSN analysis. F.X., C.F., and R.M. wrote the manuscript with input from all authors. C.F. and R.M. conceived and designed the study.

## STAR Methods

### RESOURCE AVAILABILITY

#### Lead Contact

Further information and requests for resources and reagents should be directed to and will be fulfilled by the Lead Contact, Prof. Rolf Müller.

#### Materials Availability

Plasmids generated in this study are available from the Lead Contact without restriction.

#### Data and Code Availability

The NMR spectra of synthetic substrates and their intermediates are listed in Data S1. The DNA sequences of codon-optimized *icuL* and *tgaD* are listed in Data S2. The genome sequence of strain So ce836 has been previously deposited in NCBI database with accession number of CP012672.1. The genes in BGC *icu* include IDs from SOCE836_041090 to SOCE836_041200.

#### Fermentation and HPLC-MS analysis of the icumazole producing strains

The three icumazole producers, *Sorangium cellulosum* So ce377, So ce836 and So ce950 were cultivated on M agar (10 g/L phytone, 10 g/L maltose, 1 g/L CaCl_2_, 2.047 g/L MgSO_4_ · 7 H_2_O, 11.9 g/L HEPES and 8 mg/L Fe-EDTA, 20 g/L agar, pH7.2) for 5–7 day incubation at 30 °C. Cells from each agar were collected and pre-cultured in 50 mL M liquid medium for 3–5 days at 300rpm/30 °C as seed. For fermentation, 5 mL of each seed culture was inoculated in 50 mL M liquid medium together with 1 mL pretreated XAD-16 resin and shaked at 300 rpm/30 °C. After the 8-day cultivation, cells and XAD-16 resin from each culture were harvested (8,000 rpm for 10 min) and washed with 50 mL methanol, giving rise to the crude extract.

The crude extracts were then subjected to HPLC-MS analysis with the gradient changing from 5% to 95 % B in 18.5 min under the flow rate of 0.6 mL/min. (LC: Ultimate 3000 RS; MS: Bruker Maxis II (4Generation) oq-TOF; Column: ACQUITY UPLC BEH C18 Column, 130Å, 1.7 μm, 2.1 mm X 100 mm.)

#### Bioinformatic analysis

The potential BGCs were predicted using antiSMASH 6.0 (Blin et al., 2021). The sequences of each type of domain were retrieved for further analysis. Sequence alignments of each type of gene were performed with Clustal Omega Version 1.2.2 (Baxevanis et al., 2020) embedded in Geneious Prime^®^ 2021.2.1 (Biomatters Ltd., New Zealand). The TEs of known compounds were retrieved from MIBiG 2.0 (Kautsar et al., 2020). The phylogenetic tree of TEs was built by Jukes-Cantor genetic distance model with neighbor-joining (NJ) method. The resampling method was bootstrap with 1000 replicates.

#### Molecular cloning, overexpression, and purification CCR proteins

The codon-optimized sequences of *tgaD* and *icuL* were synthesized by GenScript Biotech (Netherlands). All three CCR genes (codon-optimized *tgaD*, codon-optimized *icuL* and *leu10*) were amplified using primers and then cloned into the *Nde*I and *Hind*III pre-treated pHisTev or pHisMbpTev by NEBuilder HiFi DNA Assembly Cloning Kit (New England Biolabs Inc.). The resulting plasmids, namely pHisTev::*tgaD*, pHisMbpTev::*icuL*, pHisTev::*leu10*, were electroporated into BL21(DE3) for protein overexpression.

For protein expression (TgaD, IcuL and Leu10), a single colony was inoculated into LB liquid plus 50 μg/mL kanamycin for overnight shaking at 37 °C/180 rpm. The culture was inoculated into 6 flasks containing 1 L fresh LB (1:100) and cultured under the same condition. When the OD_600_ achieved 0.4–0.6, a final concentration of 0.1 mM isopropyl-β-D-1-thiogalactopyranoside (IPTG) was added for induction.

After overnight induction at 18 °C/200 rpm, cell mass was collected by centrifugation at 8000 rpm for 15 min at 4 °C and washed twice with lysis buffer (20 mM Tris-HCl, 200 mM NaCl, 20 mM imidazole and 10% glycerol (w/v), pH 8.0). The washed cell pellet was re-suspended in 30 mL lysis buffer and then lysed by French Press. The cell lysate was centrifuged at 23,000 rpm for 30 min at 4 °C. The supernatant was flowed through the HisTrap FF affinity column (GE), then washed by 5 column-volume of lysis buffer, and eluted by elution buffer (20 mM Tris-HCl, 200 mM NaCl, 250 mM imidazole and 10% glycerol (w/v), pH 8.0) to obtain recombinant CCR protein. The purified GetF was then desalinated and concentrated using PD-10 desalting column and Amicon, respectively.

#### *In vitro* assay

The 50 μL reaction system contains 1 mM of each substrate, 100 mM NaHCO_3_ or NH_4_HCO_3_, 4 mM NADPH or NADH, 100 mM Tris-HCl pH 8.0, and 2 μM CCR (TgaD, IcuL or Leu10). The reaction volumes without CCR served as control. The reactions were conducted at 30 °C overnight and quenched by the addition of 1% trichloroacetic acid (TCA) and 50 μL MeOH and after centrifugation at 15,000 rpm the supernatants were analyzed by HPLC-MS.

To figure out the potential reason causing the degradation of substrates, each component, namely the buffers, NaHCO_3_, enzymes, and H_2_O, was incubated with 2-octenoyl-SNAC (**S6**) overnight at 30 °C, respectively.

To explore the suitable condition for TgaD activity, various buffers (Tris-HCl buffer pH 8.0, Tris-HCl buffer pH 7.0, Phosphate potassium buffer pH 7.0, MOPS buffer pH 6.0 or MES buffer pH 6.0) or different concentrations of NaHCO_3_/NH_4_HCO_3_ (100 mM, 50 mM, 25 mM, or 0 mM) were applied for 2h incubation at 30 °C.

#### Kinetic studies of TgaD

The kinetics were performed with NADPH and NADH at gradient concentrations ranging from 0.2 mM to 2.0 mM. The reaction mixture (100 μL) contained 100 mM Tris-HCl buffer pH 7.0, 1.84 nM TgaD, 2.8 mM 2-octenoyl-SNAC, and NADPH/NADH at varying concentrations. The reactions were initiated by adding different concentrations of NADPH or NADH. The reaction was monitored spectrophotometrically at 340 nm using a 96-well plate. The measurement was performed for 15 min. The data were fitted by Origin2022. In addition, the standard curves of both NADPH and NADH were made from 0.0625 mM to 2 mM. Three biological replicates were performed. All error bars represent SEM.

#### Networking analysis of CCR genes

To construct a sequence similarity network (SSN) of CCR genes, all members of the Pfam (Mistry et al., 2021) family ADH_zinc_N (PF00107) were retrieved from UniProt (The UniProt Consortium., 2021) and aligned with HHMer (Eddy, 1998) using the Pfam family HMM along with the CCR sequences described here. After that, all sequences longer than 600 amino acids were removed, leading to 5,410 proteins predicted to lie within biosynthetic gene clusters (BGCs), which were predicted and annotated with antiSMASH (Blin et al., 2021). Sequence identity was calculated based on the alignment constructed with HMMer. Protein sequences were clustered based on the sequence identity, where all edges corresponding to sequence identity < 25% were removed. After that, all clusters comprising one single protein were removed as well. The resulting SSN was visualized with CytoScape (Shannon et al., 2003).

#### Synthesis of SNAC-esters

##### General experimental details

All air-or moisture-sensitive reactions were carried out in dried glassware (>100 °C) under an atmosphere of nitrogen or argon. Dried solvents were distilled before use. The products were purified by flash chromatography on silica gel columns (Macherey-Nagel 60, 0.04-0.063 mm). Analytical TLC was performed on pre-coated silica gel plates (Macherey-Nagel, Polygram^®^SIL G/UV254). Visualization was accomplished with UV-light, KMnO_4_ or a ceric ammonium molybdate chamber. The products were purified by flash chromatography on silica gel columns (Macherey-Nagel 60, 0.04-0.063 mm). ^1^H, and ^13^C spectra were recorded with a Bruker AV 500 [500 MHz, (^1^H), 126 MHz (^13^C)]. Spectrometer in CHCl_3_-*d* unless otherwise specified. Chemical shifts are given in parts per million (ppm) and referenced against the residual proton or carbon resonances of the >99% deuterated solvents as internal standard. Coupling constants (*J*) are given in Hertz (Hz). Data are reported as follows: chemical shift, multiplicity (s = singlet, d = doublet, t =triplet, q = quartet, quint = quintet, m = multiplet, dd = doublet of doublets, dt =doublet of triplets, br = broad and combinations of these) coupling constants, and integration. NMR spectra were evaluated using ACDLabs 2019. High-resolution mass was determined by LC-MS/MS using Thermo Scientific Q Exactive Focus Orbitrap LC-MS/MS system.

##### General procedure 1 (GP1): Wittig olefination (Nakamura et al., 2003)

A solution of aldehyde (1.0 eq.) and ethyl (triphenylphosphoranylidene)acetate (1.2 eq.) in DCM (0.5 M) was stirred at room temperature overnight. After full conversion (TLC), the solvent was removed under reduced pressure, and the crude product was purified *via* flash chromatography to give the unsaturated ester.

##### General procedure 2 (GP2): LiOH·H_2_O mediated saponification (Angelini et al., 2012)

LiOH·H_2_O (3.0 eq.1.0 M in H_2_O) was added to a solution of ethyl ester (1.0 eq.) in THF (0.2 M) at 0 °C and was stirred at room temperature overnight. After complete conversion (TLC), the solvent was removed under reduced pressure and the residue was dissolved in H_2_O before being acidified with 1 N KHSO_4_ solution to pH 1–2 at 0 °C. Extraction with DCM, drying over anhydrous Na_2_SO_4_, filtering and concentration under vacuum led to the acid without further purification.

##### General procedure 3 (GP3): SNAc-ester formation from carboxylic acid (Roberts et al., 2017)

To a solution of EDC·HCl (1.0 eq.), DIPEA (1.0 eq.), DMAP (0.1 eq.) and. carboxylic acid (1.0 eq.) in DCM (0.1 M) *N*-acetylcysteamine (1.0 eq.) was added at 0 °C. The reaction mixture was warmed up room temperature and after complete conversion (TLC) diluted with DCM and washed successively with sat. NaHCO_3_ solution, 1 N KHSO4 solution and brine. The organic phase was dried over anhydrous Na_2_SO_4_, filtered and the solvent was removed under reduced pressure. Flash chromatography afforded the desired product.

#### Ethyl (*E*)-4-methylpent-2-enoate (S1a)

According to **GP1**, isobutyraldehyde (144.2 mg, 2.00 mmol) and ethyl (triphenylphosphoranylidene)acetate (835.9 mg, 2.40 mmol) were converted into ethyl (*E*)-4-methylpent-2-enoate (**S1a**). After purification by column chromatography on silica (pentane/Et_2_O = 9: 1), unsaturated ester **S1a** (188.6 mg, 1.32 mmol, 66 %) was obtained as a colorless oil.

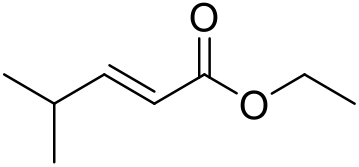

TLC: R_f_(**S1a**) = 0.29 (pentane/Et_2_O = 9:1). ^1^H NMR (500 MHz, CDCl_3_) δ = 1.07 (d, *J*=6.87 Hz, 6 H), 1.30 (t, *J*=7.17 Hz, 3 H), 2.41 – 2.51 (m, 1 H), 4.19 (q, *J*=7.07 Hz, 2 H), 5.77 (dd, *J*=15.72, 1.53 Hz, 1 H), 6.95 (dd, *J*=15.72, 6.71 Hz, 1 H). ^13^C NMR (126 MHz, CDCl_3_) δ = 14.3, 21.2, 30.9, 60.1, 118.6, 155.4, 167.1. HRMS(ESI+) *m/z* calculated for C_8_H_15_O_2_ [M+H]^+^143.1067; found, 143.1066.

#### Ethyl (*E*)-5-methylhex-2-enoate (S2a)

According to **GP1**, 3-methylbutanal (172.3 mg, 2.00 mmol) and ethyl (triphenylphosphoranylidene)acetate (835.9 mg, 2.40 mmol) were converted into ethyl (*E*)-5-methylhex-2-enoate (**S2a**). After purification by column chromatography on silica (pentane/Et_2_O = 9: 1), unsaturated ester **S2a** (299.6 mg, 1.91 mmol, 96 %) was obtained as a colorless oil.

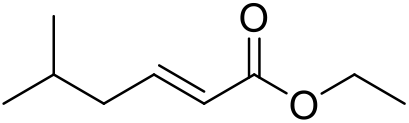

TLC: R_f_(**S2a**) = 0.30 (pentane/Et_2_O = 9:1). ^1^H NMR (500 MHz, CDCl_3_) δ = 0.93 (d, *J*=6.71 Hz, 6 H), 1.30 (t, *J*=7.10 Hz, 3 H), 1.75 – 1.82 (m, 1 H), 2.07 – 2.12 (m, 2 H), 4.19 (q, *J*=7.17 Hz, 2 H), 5.81 (dt, *J*=15.56, 1.37 Hz, 1 H), 6.95 (dt, *J*=15.11, 7.30 Hz, 1 H). ^13^C NMR (126 MHz, CDCl_3_) δ = 14.3, 22.3, 27.8, 41.4, 60.1, 122.3, 148.2, 166.7. HRMS(ESI+) *m/z* calculated for C_9_H_17_O_2_ [M+H]^+^157.1223; found, 157.1221.

#### Ethyl (*S*,*E*)-4-methylhex-2-enoate (S3a)(Nakamura et al., 2003)

According to **GP1**, (*S*)-2-methylbutanal (172.3 mg, 2.00 mmol) and ethyl (triphenylphosphoranylidene)acetate (835.9 mg, 2.40 mmol) were converted into ethyl (*S*,*E*)-4-methylhex-2-enoate (**S3a**). After purification by column chromatography on silica (pentane/Et_2_O = 9: 1), chiral unsaturated ester **S3a** (144.5 mg, 0.92 mmol, 46 %) was obtained as a colorless oil.

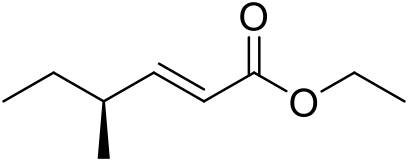

TLC: R_f_(**S3a**) = 0.35 (pentane/Et_2_O = 9:1). ^1^H NMR (500 MHz, CDCl_3_) δ = 0.89 (t, *J*=7.40 Hz, 3 H), 1.05 (d, *J*=6.71 Hz, 3 H), 1.30 (t, *J*=7.17 Hz, 3 H), 1.41 (quintd, *J*=7.27, 1.91 Hz, 2 H), 2.18 – 2.27 (m, 1 H), 4.19 (q, *J*=7.17 Hz, 2 H), 5.78 (dd, *J*=15.72, 1.07 Hz, 1 H), 6.87 (dd, *J*=15.72, 7.78 Hz, 1 H). ^13^C NMR (126 MHz, CDCl_3_) δ = 11.6, 14.3, 18.9, 28.8, 38.1, 60.2, 119.7, 154.5, 167.0. HRMS(ESI+) *m/z* calculated for C_9_H_17_O_2_ [M+H]^+^157.1223, found, 157.1222.

#### (*E*)-4-Methylpent-2-enoic acid (S1b)

According to **GP2**, ethyl (*E*)-4-methylpent-2-enoate (**S1a**) (165.5 mg, 1.16 mmol) was saponified with LiOH·H_2_O (145.7 mg, 3.47 mmol) to get the desired unsaturated acid **S1b** (129.4 mg, 1.13 mmol, 98%) as a colorless oil.

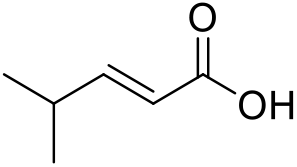

TLC: R_f_(**S1b**) = 0.13 (hexane/ethyl acetate = 9:1). ^1^H NMR (500 MHz, CDCl_3_) δ = 1.09 (d, *J*=6.87 Hz, 6 H), 2.45 – 2.55 (m, 1 H), 5.79 (dd, *J*=15.72, 1.37 Hz, 1 H), 7.07 (dd, *J*=15.72, 6.71 Hz, 1 H), 11.78 (br s, 1 H). ^13^C NMR (126 MHz, CDCl_3_) δ = 21.1, 31.1, 117.9, 158.3. HRMS(ESI+) *m/z* calculated for C_6_H_11_O_2_ [M+H]^+^115.0754; found, 115.0754.

#### (*E*)-5-Methylhex-2-enoic acid (S2b)

According to **GP2**, ethyl (*E*)-5-methylhex-2-enoate (**S2a**) (283.2 mg, 1.81 mmol) was saponified with LiOH·H_2_O (228.2 mg, 5.43 mmol) to get the desired unsaturated acid **S2b** (229.1 mg, 1.79 mmol, 99%) as a colorless oil.

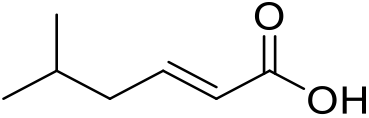

TLC: R_f_(**S2b**) = 0.16 (hexane/ethyl acetate = 9:1). ^1^H NMR (500 MHz, CDCl_3_) δ = 0.94 (d, *J*=6.71 Hz, 6 H), 1.74 – 1.95 (m, 1 H) 2.12 – 2.18 (m, 2 H), 5.83 (dt, *J*=15.56, 1.37 Hz, 1 H), 7.04 – 7.11 (m, 1 H), 11.82 (br s, 1 H). ^13^C NMR (126 MHz, CDCl_3_) δ = 22.3, 27.7, 41.5, 121.6, 151.3, 171.9. HRMS(ESI+) *m/z* calculated for C_7_H_13_O_2_ [M+H]^+^129.0910; found, 129.0908.

#### (*S*,*E*)-4-Methylhex-2-enoic acid (S3b)

According to **GP2**, ethyl (*S*,*E*)-4-methylhex-2-enoate (**S3a**) (145.0 mg, 0.93 mmol) was saponified with LiOH·H_2_O (116.84 mg, 2.78 mmol) to get the desired unsaturated acid **2c** (118.8 mg, 0.93 mmol, quant.) as a colorless oil.

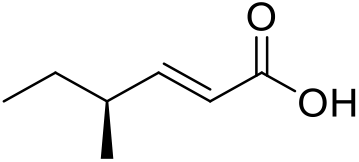

TLC: R_f_(**2c**) = 0.14 (hexane/ethyl acetate = 9:1). ^1^H NMR (500 MHz, CDCl_3_) δ = 0.90 (t, *J*=7.40 Hz, 3 H), 1.07 (d, *J*=6.71 Hz, 3 H), 1.37 – 1.49 (m, 2 H), 2.22 – 2.31 (m, 1 H), 5.80 (dd, *J*=15.72, 1.22 Hz, 1 H) 7.00 (dd, *J*=15.64, 7.86 Hz, 1 H) 11.90 (br s, 1 H). ^13^C NMR (126 MHz, CDCl_3_) δ = 11.6, 18.8, 28.7, 38.2, 119.0, 157.5, 172.3. HRMS(ESI+) *m/z* calculated for C_7_H_13_O_2_ [M+H]^+^129.0910; found, 129.0909.

#### *S*-(2-Acetamidoethyl) (*E*)-4-methylpent-2-enethioate (S1)

According to **GP3**, (*E*)-4-methylpent-2-enoic acid (**S1b**) (126.6 mg, 1.11 mmol) was reacted with EDC·HCl (212.6 mg, 1.11 mmol), DIPEA (143.3 mg, 1.11 mmol), DMAP (13.5 mg, 0.11 mmol) and *N*-acetylcysteamine (132.2 mg, 1.11 mmol) to SNAc-ester **S1** (51.9 mg, 0.24 mmol, 22%), which was obtained as a colorless oil after purification *via* flash chromatography on silica (CHCl_3_/MeOH = 95:5).

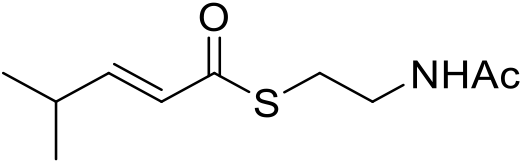

TLC: R_f_(**S1**) = 0.18 (CHCl_3_/MeOH = 95:5). ^1^H NMR (500 MHz, CDCl_3_) δ = 1.08 (d, *J*=6.87 Hz, 6 H), 1.96 (s, 3 H), 2.41 – 2.51 (m, 1 H), 3.09 (t, *J*=6.41 Hz, 2 H), 3.46 (q, *J*=6.10 Hz, 2 H), 6.01 (br s, 1 H), 6.07 (dd, *J*=15.64, 1.45 Hz, 1 H), 6.90 (dd, *J*=15.64, 6.64 Hz, 1 H). ^13^C NMR (126 MHz, CDCl_3_) δ = 21.1, 23.2, 28.2, 31.0, 39.8, 125.6, 152.5, 170.3, 190.7. HRMS(ESI+) *m/z* calculated for C_10_H1_8_NO_2_S [M+H]^+^ 216.1053; found, 216.1050.

#### *S*-(2-Acetamidoethyl) (*E*)-5-methylhex-2-enethioate (S2)

According to **GP3**, (*E*)-5-methylhex-2-enoic acid (**S2b**) (225.3 mg, 1.76 mmol) was reacted with EDC·HCl (336.9 mg, 1.76 mmol), DIPEA (227.0 mg, 1.76 mmol), DMAP (21.5 mg, 0.18 mmol) and *N*-acetylcysteamine (209.5 mg, 1.76 mmol) to SNAc-ester **S2** (107.9 mg, 0.47 mmol, 27%), which was obtained as a colorless oil after purification *via* flash chromatography on silica (CHCl_3_/MeOH = 95:5).

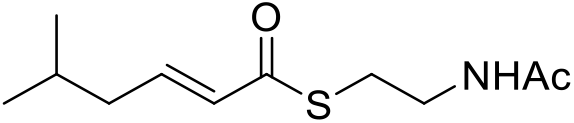

TLC: R_f_(**S2**) = 0.30 (CHCl_3_/MeOH = 95:5). ^1^H NMR (500 MHz, CDCl_3_) δ = 0.94 (d, *J*=6.71 Hz, 6 H), 1.73 – 1.85 (m, 1 H), 1.95 – 1.99 (m, 3 H), 2.09 – 2.13 (m, 2 H), 3.10 (t, *J*=6.33 Hz, 2 H), 3.48 (q, *J*=5.95 Hz, 2 H), 5.89 (br s, 1 H), 6.13 (dt, *J*=15.56, 1.37 Hz, 1 H), 6.88 – 6.96 (m, 1 H). ^13^C NMR (126 MHz, CDCl_3_) δ = 22.4, 23.2, 27.8, 28.3, 39.9, 41.5, 129.3, 145.6, 170.2, 190.6. HRMS(ESI+) *m/z* calculated for C_11_H_20_NO_2_S [M+H]^+^ 230.1209; found, 230.1205.

#### *S*-(2-Acetamidoethyl) (*S*,*E*)-4-methylhex-2-enethioate (S3)

According to **GP3**, (*S*,*E*)-4-methylhex-2-enoic acid (**S3b**) (110.0 mg, 0.86 mmol) was reacted with EDC·HCl (181.0 mg, 0.94 mmol), DIPEA (122.0 mg, 0.94 mmol), DMAP (10.5 mg, 0.09 mmol) and *N*-acetylcysteamine (112.5 mg, 0.94 mmol) to SNAc-ester **S3** (75.4 mg, 0.32 mmol, 38%), which was obtained as a colorless oil after purification *via* flash chromatography on silica (CHCl_3_/MeOH = 95: 5).

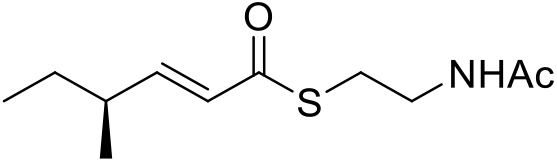

TLC: R_f_(**S3**) = 0.40 (CHCl_3_/MeOH = 95:5). ^1^H NMR (500 MHz, CDCl_3_) δ = 0.89 (t, *J*=7.48 Hz, 3 H), 1.06 (d, *J*=6.71 Hz, 3 H), 1.43 (quintd, *J*=7.27, 1.60 Hz, 2 H), 1.98 (s, 3 H), 2.19 – 2.27 (m, 1 H), 3.10 (t, *J*=6.41 Hz, 2 H), 3.48 (q, *J*=6.00 Hz, 2 H), 5.90 – 6.01 (m, 1 H), 6.10 (dd, *J*=15.64, 1.14 Hz, 1 H), 6.84 (dd, *J*=15.56, 7.78 Hz, 1 H). ^13^C NMR (126 MHz, CDCl_3_) δ = 11.6, 18.8, 23.2, 28.2, 28.7, 28.7, 38.2, 39.9, 126.7, 151.7, 170.4, 190.7. HRMS(ESI+) *m/z* calculated for C_11_H_20_NO_2_S [M+H]^+^ 230.1209; found, 230.1206.

#### *S*-(2-Acetamidoethyl) (*E*)-pent-2-enethioate (S4)

According to **GP3**, commercially available (*E*)-pent-2-enoic acid (220.2 mg, 2.00 mmol) was reacted with EDC·HCl (383.4 mg, 2.00 mmol), DIPEA (258.5 mg, 2.00 mmol), DMAP (24.4 mg, 0.20 mmol) and *N*-acetylcysteamine (238.4 mg, 2.00 mmol) to SNAc-ester **S4** (120.0 mg, 0.60 mmol, 30%), which was obtained as a colorless oil after purification *via* flash chromatography on silica (DCM/MeOH = 95:5).

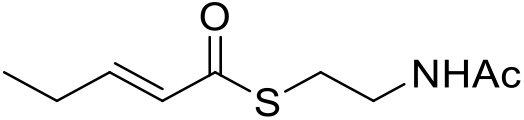

TLC: R_f_(**S4**) = 0.40 (DCM/MeOH = 9:1). ^1^H NMR (500 MHz, CDCl_3_) δ = 1.10 (t, *J*=7.48 Hz, 3 H), 1.97 (s, 3 H), 2.20 – 2.29 (m, 2 H), 2.95 – 3.12 (m, 2 H), 3.31 – 3.52 (m, 2 H), 5.89 (br s, 1 H), 6.14 (dt, *J*=15.53, 1.62 Hz, 1 H), 7.00 (dt, *J*=15.56, 6.33 Hz, 1 H).^13^C NMR (126 MHz, CDCl_3_) δ ppm 12.0, 23.3, 25.3, 28.2, 39.8, 127.4, 148.0, 170.2, 190.5. HRMS(ESI+) *m/z* calculated for C_9_H_16_NO_2_S [M+H]^+^ 202.0896; found, 202.0895.

#### *S*-(2-Acetamidoethyl) (*E*)-hex-2-enethioate (S5)

According to **GP3**, commercially available (*E*)-hex-2-enoic acid (228.3 mg, 2.00 mmol) was reacted with EDC·HCl (383.4 mg, 2.00 mmol), DIPEA (258.5 mg, 2.00 mmol), DMAP (24.4 mg, 0.20 mmol) and *N*-acetylcysteamine (238.4 mg, 2.00 mmol) to SNAc-ester **S5** (93.5 mg, 0.43 mmol, 22%), which was obtained as a colorless oil after purification *via* flash chromatography on silica (DCM/MeOH = 95:5).

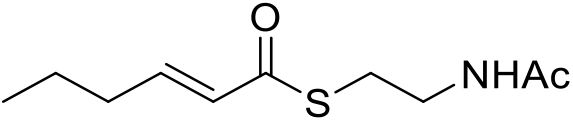

TLC: R_f_(**S5**) = 0.50 (DCM/MeOH = 9:1). ^1^H NMR (500 MHz, CDCl_3_) δ = 0.93 (t, *J*=7.32 Hz, 3 H), 1.49 (sxt, *J*=7.39 Hz, 2 H), 1.95 (s, 3 H), 2.17 (qd, *J*=7.22, 1.53 Hz, 2 H), 3.07 (t, *J*=6.48 Hz, 2 H), 3.44 (q, *J*=6.21 Hz, 2 H), 6.04 – 6.21 (m, 2 H), 6.91 (dt, *J*=15.53, 6.96 Hz, 1 H). ^13^C NMR (126 MHz, CDCl_3_) δ = 13.6, 21.1, 23.1, 28.2, 34.1, 39.7, 128.4, 146.4, 170.3, 190.3. HRMS(ESI+) *m/z* calculated for C_10_H_18_NO_2_S [M+H]^+^ 216.1053; found, 216.1052.

#### *S*-(2-Acetamidoethyl) (*E*)-oct-2-enethioate (S6)

According to **GP3**, commercially available (*E*)-oct-2-enoic acid (284.2 mg, 2.00 mmol) was reacted with EDC·HCl (383.4 mg, 2.00 mmol), DIPEA (258.5 mg, 2.00 mmol), DMAP (24.4 mg, 0.20 mmol) and *N*-acetylcysteamine (238.4 mg, 2.00 mmol) to SNAc-ester **S6** (224.7 mg, 0.92 mmol, 46%), which was obtained as a colorless oil after purification *via* flash chromatography on silica (DCM/MeOH = 95:5).

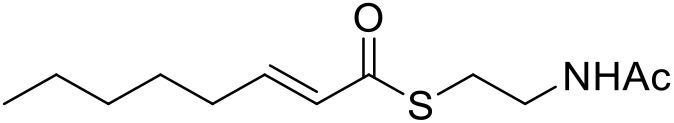

TLC: R_f_(**S6**) = 0.50 (DCM/MeOH = 9:1). ^1^H NMR (500 MHz, CDCl_3_) δ = 0.90 (t, *J*=7.05 Hz, 3 H), 1.24 – 1.39 (m, 4 H), 1.44 – 1.50 (m, 2 H), 1.97 (s, 3 H), 2.21 (qd, *J*=7.29, 1.47 Hz, 2 H), 3.10 (t, *J*=6.35 Hz, 2 H), 3.47 (q, *J*=5.94 Hz, 2 H), 5.83 – 5.98 (m, 1 H), 6.13 (dt, *J*=15.52, 1.47 Hz, 1 H), 6.94 (dt, *J*=15.52, 6.99 Hz, 1 H). ^13^C NMR (126 MHz, CDCl_3_) δ = 13.9, 22.4, 23.2, 27.5, 28.2, 31.3, 32.2, 39.8, 128.2, 146.9, 170.3, 190.5. HRMS(ESI+) *m/z* calculated for C_12_H_22_NO_2_S [M+H]^+^ 244.1366; found, 244.1362.

